# Homo-harringtonine (HHT) – A highly effective drug against coronaviruses and the potential for large-scale clinical applications

**DOI:** 10.1101/2021.04.16.440104

**Authors:** Hai-Jun Wen, Pei Lin, Gong-Xun Zhong, Zhi-Chao Xu, Lei Shuai, Zhi-Yuan Wen, Chong Wang, Xue Cao, Wen-Bin He, Jing Feng, Qi-Chun Cai, Hua-Juan Ma, Si-Jin Wu, Guo-Dong Wang, Xue-Mei Lyu, Feng-Liang Liu, Yong-Tang Zheng, Hui Zeng, Xiong-Lei He, Hualan Chen, Fu-Jie Zhang, Chung-I Wu

## Abstract

In the search for treatment schemes of COVID-19, we start by examining the general weakness of coronaviruses and then identify approved drugs attacking that weakness. The approach, if successful, should identify drugs with a specific mechanism that is at least as effective as the best drugs proposed and are ready for clinical trials. All coronaviruses translate their non-structural proteins (∼16) in concatenation, resulting in a very large super-protein. Homo-harringtonine (HHT), which has been approved for the treatment of leukemia, blocks protein elongation very effectively. Hence, HHT can repress the replication of many coronaviruses at the nano-molar concentration. In two mouse models, HHT clears SARS-CoV-2 in 3 days, especially by nasal dripping of 40 ug per day. We also use dogs to confirm the safety of HHT delivered by nebulization. The nebulization scheme could be ready for large-scale applications at the onset of the next epidemics. For the current COVID-19, a clinical trial has been approved by the Ditan hospital of Beijing but could not be implemented for want of patients. The protocol is available to qualified medical facilities.

## Introduction

Coronaviruses have increasingly become the causal agent of epidemics in humans as well as in domesticated animals(*1, 2*). Since the beginning of this century, there have been 3 such epidemics, SARS (2003), MERS (2012) and COVID-19, and they may not be the last ones (*1, 3*). At present, there is no highly effective and broadly applicable treatment scheme to thwart the COVID-19 pandemic. Obviously, an effective treatment scheme should be desirable. Although such a scheme may or may not be in time for this current pandemic, it can certainly be ready as the first-line defense at the onset of the next coronavirus epidemics.

This current report differs from other proposed schemes for treating COVID-19 (see the next section) in that we start by identifying the general weakness of the coronaviruses and then search for approved drugs that target the viruses by a specific mechanism. In contrast, the mainstream approach is to screen for effective drugs and then identify the repression mechanisms. There is a three-fold advantage to our approach if (and only if) the weakness of the viruses and the drug targeting this weakness can be identified. First, many of the experiments would have been done and published. Second, the efficacy of the drug can be set at a level higher than the most effective agents proposed to treat the infection. Third, since the scheme would target coronaviruses in general, it should be ready at the onset of the next epidemics caused by coronaviruses.

### A brief survey of drugs proposed for treating COVID-19

To have an overview of the proposals for treating COVID-19, we carry out a survey of drugs targeting the virus itself. The survey is of a limited scope as schemes that boost or suppress the immunity against the virus (such as cytokine storms) are not included. In particular, we focus on the schemes that target the cellular components involved in the replication of SARS-CoV-2. Presumably, direct attacks on the virus itself may select strongly for viral evasion whereas the attack on the host pathways is less likely to elicit such responses(*4*). In this survey, we classify the studies by the life-cycle stage where the drug imposes its effects. The stages and publications are: attachment and entry(*5–9*), translation of polyproteins(*10–13*), proteolytic processing(*14, 15*), transcription and replication(*16–18*)and multiple stages(*19, 20*). It is generally accepted that comprehensive clinical benefits have not yet been demonstrated.

**Supplementary Table 1** summarize the results of the survey. Given the complexities and variations of these experimental studies(*5–23*), we attempt to identify proposals that fulfill the following two criteria: 1) The application of the drug to the animal model results in the clearance of SARS-CoV-2 in > 80% of the animals within a reasonable time (< one week). The clearance as shown by both the viral RNA measurement and the viral titer must be significantly faster than the control; and 2) The animals show no adverse effects at the end of the experiment. (In studies where the comparison is also made against remdesivir, we further require the performance to be no worse than that of remdesivir.)

In the summary of **Supplementary Table 1**, we did not find a published scheme that fulfills both criteria. We therefore aim to find a drug that can do so. Furthermore, it would be better if the drug is FDA-approved, readily available, inexpensive and amenable to large-scale application.

### The general weakness of coronaviruses

Before we identify a suitable drug, we shall first examine the weaknesses of the viruses. Coronaviruses possess the largest genomes (26.4 to 31.7 kb) among all known RNA viruses and use two thirds of their genomes to make a super-peptide comprising ∼16 non-structural proteins (NSPs), synthesized in concatenation and later proteolyzed into its component proteins (*24*). In SARS-CoV-2 and its relatives, the super-protein is > 700 kilodaltons and ∼8000 amino acids in size. Since human peptides are rarely larger than 5000 amino-acids long and the few large ones are often highly tissue-specific with a relatively long half-life, the viral super-protein stands out among the host’s proteins.

The peculiar way of making a super-protein for later proteolysis has been hypothesized to be the Achilles heel of SARS-CoV-2(*25*). We may extend their arguments in two ways. First, a drug that blocks the initiation of the translation can be highly effective against the virus since one blockage can abrogate all 16 NSPs. Second, if a drug works to block the elongation, it would be possible to fine-tune the dosage to block peptide translation of > 5000 amino acids, but much less so for smaller peptides.

In addition to the protein size, it has been pointed out that a common weakness of virus-infected cells is hyper-transcription or hyper-translation activities. After all, viruses often replicate at an exceedingly high rate. The heightened activities are true in cancer cells as well (*25*) and that may be why many anti-cancer and anti-virus drugs often target the transcription or translation machinery(*26*). It is hence possible that some drugs may be effective against both cancers and viruses. Generally, virus-infected cells divert up to 80% of the translation capacity to serve the unrelenting demand of the virus. The mechanism by which SARS-CoV-2 drives the heightened translation has recently been clarified(*27*).

### The specific mechanism of suppression by Homo-harringtonine (HHT)

Homo-harringtonine (HHT), or omacetaxine mepesuccinate in its semi-synthetic form, is a cytotoxic plant alkaloid extracted from *Cephalotaxus* species and is likely the strongest inhibitor of protein translation approved for clinical use(*28*). HHT has been commonly used in China to treat cancer patients since the 70’s. It is the first agent approved by FDA (USA, in 2012) targeting the mRNA translation process(*29*). Our survey(*5–23*)also found other drugs that target the cell’s translation machinery for treating COVID-19(*10–12*), including Plitidepsin(*10*).

Given its wide use, the molecular mechanism of HHT has been well understood(*30–32*). HHT is described as a drug against peptide elongation (see Fig. 1A), especially “initial elongation”(*30, 31*). The latter may simply mean a highly efficient blockage at a high (and non-clinical) dose in vitro as HHT does not affect translation initiation. Based on the structural data of Garreau de Loubresse *et al*.(*31*), Fig. 1B shows graphically how HHT competes with the amino acid side chains of aminoacyl-tRNAs for binding to the A-site cleft of the ribosome.

**Figure 1:**
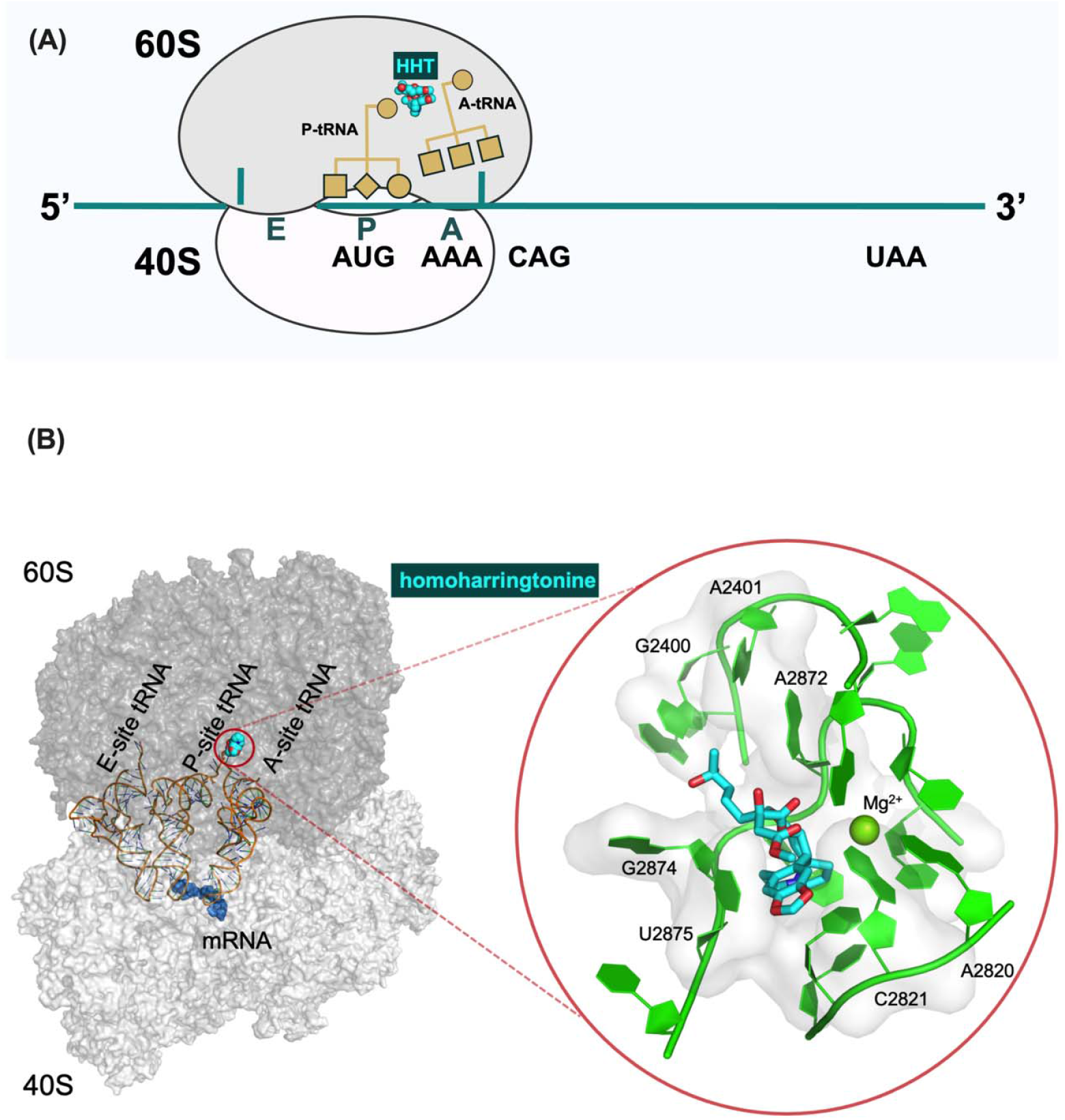
The mechanism of HHT blockage of translation elongation, which underlies its anti-coronavirus efficacy. (A) HHT prevents the incoming aminoacyl-tRNA (A-tRNA) from unloading its amino acid cargo (P-tRNA) to extend the peptide by one amino acid; (B) Structural detail of the docking of HHT at the A site of the ribosome, resulting in the inhibition of translation elongation shown in 1A. The drawing is based on the structural data (PDB code: 4U4Q) in Garreau de Loubresse *et al*. (*31*).

Thanks to the highly specific mechanism of viral repression, the HHT scheme is expected to be effective at a very low dose. The efficacy, if true, would have the following advantage in the clinical application: if we target organs with the highest viral loads (say, the respiratory system) and deliver a low dose of HHT locally (by nebulization, for example), the treatment could be effective with minimal adverse effects. The toxicity is minimized due to the low drug concentration outside of the target area.

### The efficacy of HHT against coronaviruses

Interestingly, since the SARS of 2003, there have been many efforts to identify drugs capable of repressing coronaviral replication. In each of these drug-screen studies(*33–37*), multiple drugs are reported to be effective against some coronaviruses. Curiously, there was not much emphasis that HHT (and only HHT) appears in almost all lists. Before 2020, five coronaviruses have been documented to be repressed by HHT(*33–35*)-MHV (murine coronavirus mouse hepatitis virus), Bovine coronavirus strain L9 (BCoV-L9), human enteric coronavirus strain 4408 (HECoV-4408), porcine epidemic diarrhea virus (PEDV) and MERS (**Supplementary Table 2**).

Because these reports are spread among laboratories using various dosages, the HHT effect might be due to diverse factors. To test the hypothesis of a common mechanism, we use PEDV from the previous list (*34*) and two additional porcine coronaviruses – SADS-CoV and PD-CoV in the same experimental setting. Our goal is to find out whether the dose-response among coronaviruses is indeed similarly low. Fig. 2 shows that the IC50 is generally around 100 nM and the eradication is achieved at < 1uM. We note that these are the same range of values reported in the literature(*13, 33–37*). In summary, eight different coronaviruses, including SARS-CoV-2, have been shown to be repressible by HHT at a comparable concentration. HHT thus appears to act against a general feature in protein translation.

**Figure 2:**
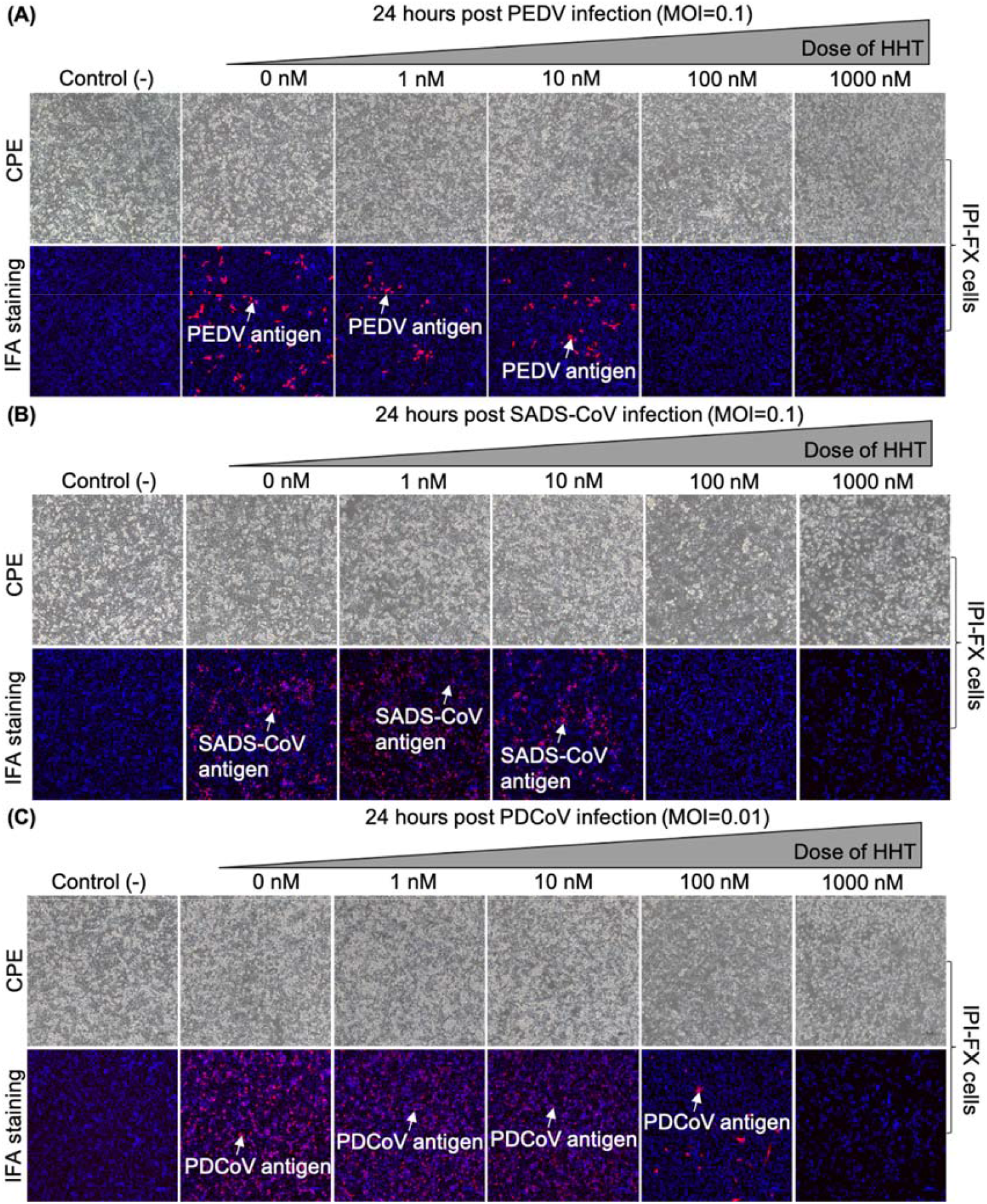
HHT inhibits 3 different porcine coronaviruses in vitro at comparable doses (< 100 nM). While coronaviruses have been reported to be susceptible to HHT inhibition (*33–37*, and **Supplementary Table 2**), the test on additional coronaviruses here is to show comparable doses in the viral inhibition in the same experimental setting. If a general mechanism exists, comparable doses should be observed.

### The efficacy of HHT in suppressing SARS-CoV-2 in mouse models

We now present data on HHT efficacy against coronaviruses in vivo. The main experiments on infected mice are done in a P4 laboratory which uses a particular SARS-CoV-2 strain capable of infecting wildtype mice(*38*). HHT is delivered via intraperitoneal injection (IP), intranasal dripping (IN) or both (IP+IN). The IP part of the experiment was replicated in another P3 laboratory that uses the transgenic mice carrying the human ACE-2 receptor (*39*) and a common SARS-CoV-2 strain.

In the initial experiments, the main question was the efficacy of viral repression, leaving the toxicity issue for later resolution. Since 1 uM concentration of HHT for 2 days can eradicate SARS-CoV-2 in vitro, we administered the drug at the theoretical concentration of 5 uM on the first day and 2.5 uM on each subsequent day, or roughly 80 ug and 40 ug per mouse per day (see Methods for the rationale). In Fig. 3A and 3B, HHT via IP infection appears highly effective in 3-4 days by either protocol. The viral load is not detectable in all but one case where the load is reduced to < 1%. Hence, viral eradication is achievable in vivo as in the in vitro assays.

**Figure 3:**
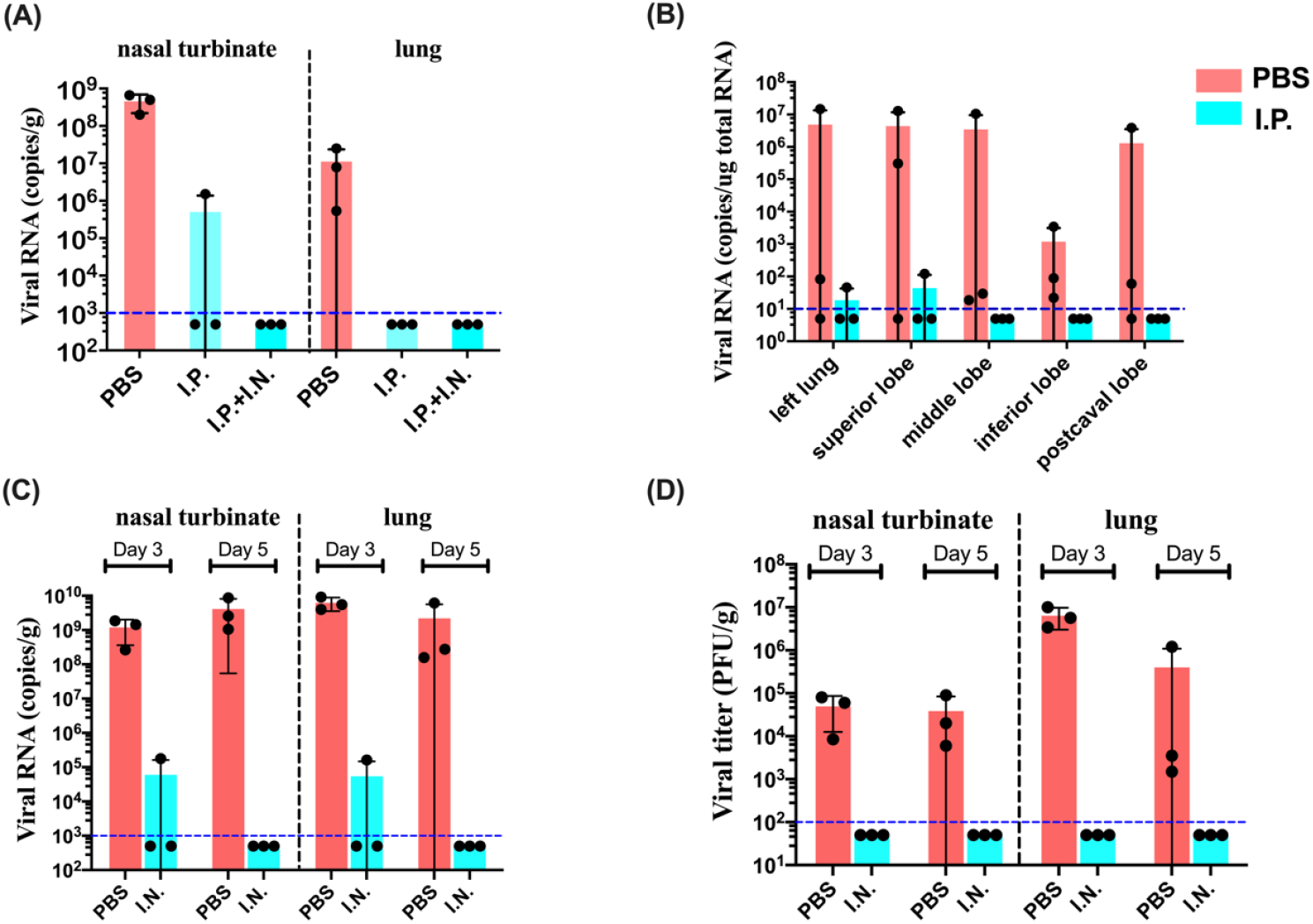
The repression of SARS-CoV-2 by HHT in mouse models. (A) I.P. (Intraperitoneal injection) or I.P.+I.N. (intranasal dripping) carried out in a P4 laboratory. The tissues assayed on the 4^th^ day after injection are given on the top. (B) The repeats of the I.P. experiment of (A) in another laboratory that uses a different mouse model and a different viral strain. The tissues assayed on the third day after injection are shown at the bottom. (C-D) I.N. only and the repression appears complete. Viral RNA and viral titer assays in the same P4 laboratory are presented. The horizontal dotted line (blue color) shows the detection limit of viral load in each assay. See the main text for details.

At the high concentration of > 1uM, the efficacy as well as some adverse effects may both be expected. We interpret 5 uM to be close to the maximal tolerated dose (MTD) for HHT in mice because even the slight variation in experimental conditions at this dose can result in fairly different outcomes (little toxicity in the experiments of Fig. 3B but substantial adverse effects in 3A). It is also interesting that a high dose near MTD for a short duration as in Fig. 3B could be effective with acceptable toxicity.

We then attempt to maintain the efficacy but avoid the toxicity. In a modified experiment that delivers the drug by nasal dripping only, the total dose is half of that in the IP+IN set. Most significantly, the efficacy in the IN experiment is the same as in the IP+IN experiment (Fig. 3A vs. Fig. 3C-3D) at half the total dose and, importantly, *sans* the adverse effects. In short, nasal dripping alone is fully effective in clearing the virus. The additional IP injection may be harmful without benefits.

### The safety dosage of HHT by nebulization

While nasal dripping works in mice, it may not work in larger animals. We therefore carry out nebulization on dogs to measure the toxicity in larger mammals. At 10-12 kg, dogs should be more comparable with humans in toxicity tolerance although they do not get infected by SARS-CoV-2(*40*). Among animal models, dogs are exceptional in being able to cooperate to receive daily nebulization without anesthetization. The particles in nebulization, ∼ 5 um in diameter, are expected to be able to reach the lungs. In our experiment, the dosage is 0 mg, 0.5 mg, 1 mg and 1.5 mg per day for 7-10 days. By the end of the experiment, all dogs appear normal in body weight, blood cell count and blood biochemistry as shown in Fig. 4A-C. The dog receiving the highest dose was euthanized and the autopsy appears normal as well. The remaining dogs have been doing well. In short, dogs can tolerate HHT nebulization at doses much higher than the calculated dosage for treating human patients.

**Figure 4:**
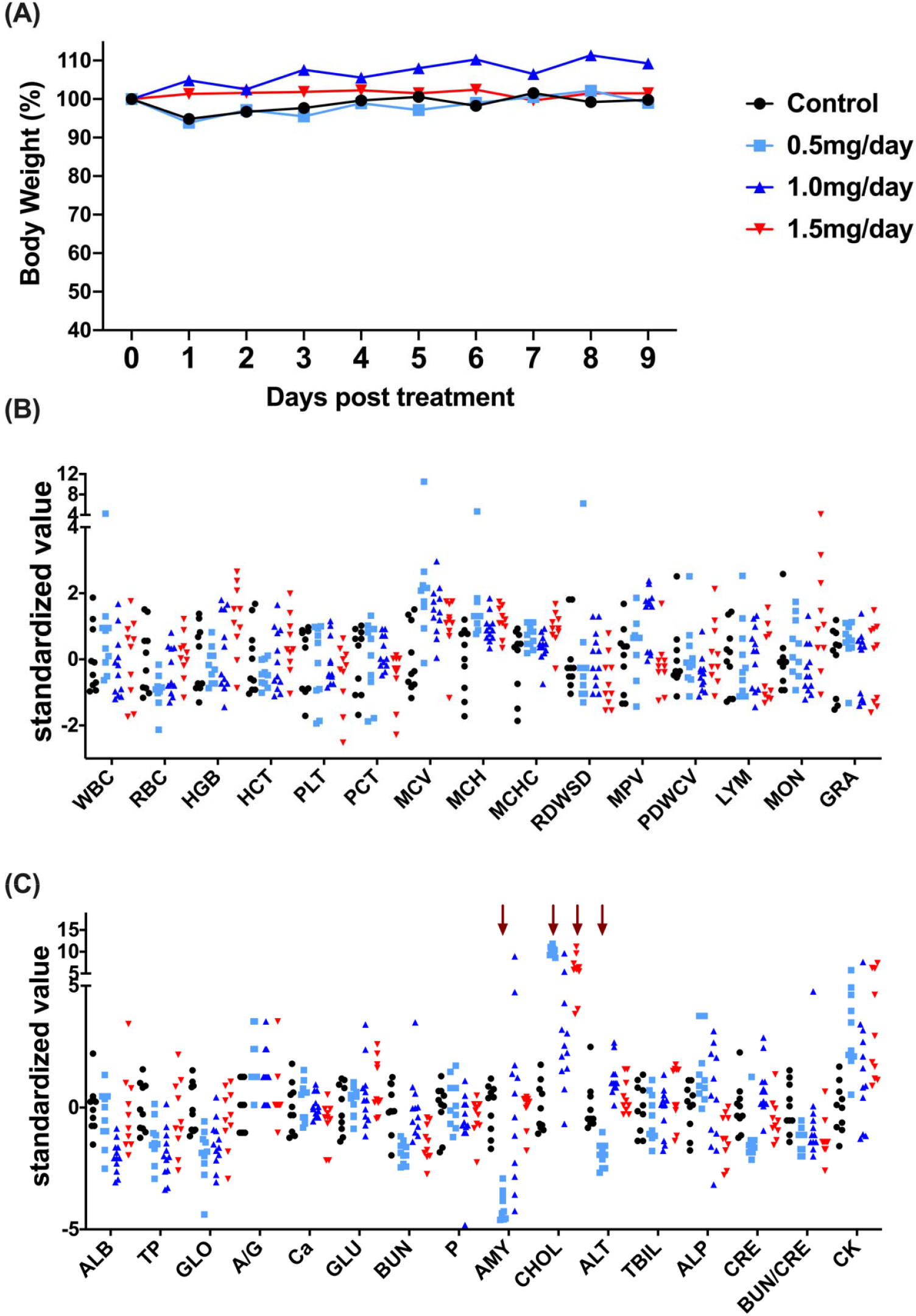
Evaluation the safety of HHT delivered by nebulization in a canine model. (A) Body weight changes during treatment. No adverse effects were detected. (B) complete blood cell count. (C) Biochemistry test. Values at (B) and (C) were standardized by z_i,j_ = (x_i,j_ - μ_j_) / σ_j_ of the normal distribution. Each dot is a sample, taken every day, from an experimental dog. For each test, three treated groups were compared with the control separately using Mann-Whitney U test and Bonferroni correction for multiple comparisons. The four measurements (amylase (AMY), cholesterol (CHOL), and glutamic-pyruvic transaminase (ALT) in the 0.5mg/day group, and CHOL in the 1.5mg/day group) which were significantly different (p<0.01 after correction for multiple comparisons) from the value in control group were marked by mocha arrows. Since all values in the four measurements were within the reference range, we concluded that these dosages administrated by nebulization were well-tolerated in dogs.

## Discussion

HHT is effective in repressing all 8 coronaviruses tested in vitro(*13, 33–37*). The drug concentration necessary for viral clearance in vitro is consistently under 1 uM. (An anomaly in the literature is Choy *et al*. (2020)(*41*) which shows discrepancies with the literature in multiple drugs; see Wen *et al*. (2021)(*13*) and Ianevski *et al*. (2020)(*37*)) As stated above, the specific mechanism of HHT repression of viral protein translation may be the key.

For the treatment scheme to be ready for the next coronavirus epidemic in either humans or animals, an unresolved issue is the mode of delivering HHT. Although intravenous injection (I.V.) is the standard delivery method to treat leukemia and the safety level is well known(*29, 42*), I.V. is not necessarily the best option for treatment. I.P. injection in mice shows evidence of toxicity in one of the two experiments (Fig. 3A vs. 3B). Overall, the systemic administration is neither desirable nor necessary if the goal is to reduce the viral burden with minimal toxicity.

For the potential clinical applications of HHT on coronavirus patients, animal experiments point to nebulization. By nebulization, the drug is concentrated in the respiratory track, where the viral load is the highest(*43*). While the virus is known to invade other organs, the clearance from the respiratory track may allow the immune system to cope with the remaining viral loads. Using a portable device (like the one used for asthma), nebulization can be used on a large number of infected patients without hospital stay.

With the rapid developments of vaccines, the HHT treatment scheme should certainly be thought of as one for the future. As for the current epidemics, we have developed a protocol for a clinical trial in the Ditan hospital where all COVID-19 patients from the Beijing area were treated. In this protocol, we consider the merger of phase I and phase II trials. Given the rate of self-recovery with COVID-19 at > 95%, most of the infections with no or light symptoms would be like healthy individuals recruited into the standard phase I trial. However, unlike healthy individuals in a phase I trial, a large number of infections with mild symptoms might benefit from the HHT application while providing the safety data, starting at the lowest doses of the standard phase I trial.

Since the protocol was approved by the IRB of Ditan in February of 2021, there have been no COVID-19 patients in or near Beijing. Until the vaccinations take full effects globally, many regions may still benefit from the HHT treatment scheme. We are therefore offering the protocol to certified medical facilities that might benefit from a clinical trial for treating COVID-19.

## Methods

### Facility, Ethics, and Biosafety statement

Main experiments with infectious SARS-CoV-2 were performed in the biosafety level 4 and animal biosafety level 4 facilities in the Harbin Veterinary Research Institute (HVRI) of the Chinese Academy of Agricultural Sciences (CAAS), approved by the Ministry of Agriculture and Rural Affairs of China.

Part of the in vivo antiviral studies were carried out at biosafety level-3 (BSL3) conditions at the Key Laboratory of Animal Models and Human Disease Mechanisms of the Chinese Academy of Sciences, Kunming Institute of Zoology (KIZ). The animal studies were carried out in strict accordance with the recommendations in the Guide for the Care and Use of Laboratory Animals of the Ministry of Science and Technology of the People’s Republic of China. The protocols were approved by the Committee on the Ethics of Animal Experiments of the HVRI of CAAS or Institutional Committee for Animal Care and Biosafety at Kunming Institute of Zoology, Chinese Academy of Sciences, respectively. All animals used in this study were chosen randomly.

### Cells and Viruses at HVRI in Harbin

Vero E6 cells were maintained in Dulbecco’s modified Eagle’s medium (DMEM) containing 10% fetal bovine serum (FBS) and antibiotics and incubated at 37°C with 5% CO2. Mouse-adapted SARS-CoV-2/HRB26/human/2020/CHN (HRB26M, GISAID access no. EPI_ISL_459910) was obtained by serially passaging the HRB26 virus in 4–6-week-old female mice until passage 14 and was propagated in Vero E6 cells. Infectious virus titers were determined by using a plaque forming unit (PFU) assay in Vero E6 cells.

### Cells and Viruses at KIZ in Kunming

The SARS-CoV-2 (strain 107) was provided by Guangdong Provincial Center for Disease Control and Prevention (Guangzhou, China). This virus was propagated and titrated on African green monkey kidney epithelial cells (Vero E6) (ATCC, no. 1586), which were cultured in Dulbecco’s modified Eagle’s medium (DMEM, Gibco) with 4.5 mM L-glutamine (GE Life Sciences) supplemented with 10% FBS (Hyclone) and 1% penicillin–streptavidin (Gibco). Cells were cultured at 37°C in a humidified 5% CO2 atmosphere. Mycoplasma testing was performed at regular intervals and no mycoplasma contamination was detected.

### *In vivo* antiviral studies of HHT at HVRI in Harbin

The mice for this study, 6-week-old female BALB/c, were obtained from Beijing Charles River Labs (Beijing, China). Mice were lightly anesthetized with CO2 and intranasally (I.N.) inoculated with 50 μL dilutions of SARS-CoV-2. Body weights and clinical symptoms were monitored daily. Groups of six mice were treated by intraperitoneal (I.P.) injection with a loading dose of 80ug/mouse HHT, followed by a daily maintenance dose of 40ug/mouse. In an allternative mode, mice were treated I.N. (40ug/mouse daily) alone or a combination of I.N. (40ug/mouse daily) and I.P. (40ug/mouse daily). As a control, mice were administered vehicle solution (PBS) daily. One hour after administration of the loading dose of HHT or vehicle solution, each mouse was inoculated i.n. with10^3.1^ PFU of HRB26M in 50μL. Three mice from each group were euthanized on days 3 and 5 post-inoculation. The nasal turbinates and lungs were collected for virus detection by qPCR and PFU assay(*38, 44*).

### *In vivo* antiviral studies of HHT at KIZ in Kunming

Experiments in KIZ were part of the multi-laboratory design but in a smaller scale than those in HVRI. The angiotensin-converting enzyme 2 (ACE2) humanized mice (hACE2 mice) aged 8-10 weeks were generated from Guangzhou Institute of Biomedicine and Health (GIBH), Chinese Academy of Sciences (CAS)(*39*). Mice were anesthetized with isoflurane (RWD Life Science, Shenzhen) and intranasally infected with 2×10^6^ TCID_50_ of SARS-CoV-2 (strain 107) in 30μl of DMEM. Body weights and clinical symptoms were monitored daily. Treated group of three mice were treated intraperitoneal (I.P.) with a loading dose of 80ug/mouse HHT, followed by a daily maintenance dose of 40ug/mouse. As a control, mice (n=3) were administered vehicle solution (PBS) daily. Lung tissues were collected on day 3 post-inoculation. RNA was extracted from lung tissues using the TRIzol™ Reagent (Invitrogen) according to the manufacturer’s instructions. Viral RNA was quantified by THUNDERBIRD® Probe One-step qRT-PCR Kit (Toyobo) according to the manufacturer’s instructions and the TaqMan primers (Forward primer: 5’-GGGGAACTTCTCCTGCTAGAAT-3’; Reverse primer: 5’-CAGACATTTTGCTCTCAAGCTG-3’. The TaqMan probe sequences were 5’-FAM-TTGCTGCTGCTTGACAGATT-TAMRA-3’.). The results were expressed as copies per microgram tissue.

### Evaluation of the anti-viral efficacy of HHT against other coronaviruses

African green monkey kidney (Vero) cells were obtained from ATCC (ATCC number: CCL-81) (USA) and Porcine intestinal epithelial cell clone J2 (IPEC-J2) cells were obtained from Wen’ s Foodstuffs Group Co, Ltd (Guangdong, China). All cells were cultured in Dulbecco’ s modified eagle medium (DMEM) (Hyclone, USA) supplemented with 100 U/mL penicillin, 100 U/mL streptomycin, and 10% fetal bovine serum (FBS) (BOVOGEN, Australia). The maintenance medium for PEDV, SADS-CoV, or PDCoV propagation was DMEM supplemented with 7.5 μg/mL trypsin (Gibco, USA)(*45*).

Confluent Vero or IPEC-J2 cell monolayers in 12-well plate were inoculated with various concentrations of HHT (1-1000nM) or the control normal DMEM for 1 h, followed by infection with PEDV, SADS-CoV, or PDCoV at an MOI of 0.1 or 0.01 for 1 h, and then the viral inoculums was removed and fresh maintenance medium containing different concentrations of HHT was added. Twenty-four hours later, cells were checked under microscopy to observe cytopathic effect (CPE) and then fixed for indirect immunofluorescent assay (IFA). Briefly, cells were fixed with 4% paraformaldehyde for 15 min and then permeabilized with 0.2% Triton X-100 for 15 min at room temperature. After blocked with 1% bovine serum albumin (BSA), cells were stained with anti-PEDV (SADS-CoV, or PDCoV) N polyclonal antibody (Wen’ s Foodstuffs Group Co., Ltd, China) (1:1000) at 37°C for 1 h. Cells were then washed with 1 × PBS and incubated with fluoresceinisothiocyanate (FITC) (1:500) or Cy3-labeled goat anti-mouse secondary antibody (KPL, USA) (1:500) at 37°C for 1 h. After three washes in 1 × PBS, cells were counter-stained with DAPI and observed with a fluorescence microscope (LEICA DMi8, Germany).

### Evaluation the safety of HHT delivered by nebulization in a canine model

Four beagles (female) weighing 10–12 kg were used in this study. The dogs were judged to be in good health based on the results of physical examinations, complete blood cell counts, and serum biochemical analyses. Each dog was fed with an appropriate amount of food and their health status was monitored daily by a dedicated veterinarian.

One dog was assigned to each of the following nebulization dosage: 0.5mg/day, 1.0mg/day, 1.5mg/day, or normal saline alone (control group). Dogs were treated for 7-10 days. Nebulization of HHT was performed in the animal research facility at the University of Kunming Medical University, using a commercially available ultrasonic nebulizer (particle size arounds 5 microns) connected to a polyethylene rebreathing bag. The polyethylene bag was held manually over the muzzle of the dog during treatments (15-20 minutes).

Blood samples were collected daily for complete blood cell counts and serum biochemical analyses. At the end of treatment, the dog received 1.5mg HHT per day was sacrificed and the lung tissue was collected for histopathological study. The other three dogs were adopted.

---

**Supplementary Table 1.** In vivo anti-SARS-CoV-2 efficacy of compounds targeting the viral life cycle.

| virus life cycle stage | Compound name | Primary indication | Clinical Phase for COVID-19 | In vivo performance (compared to either vehicle or Remdesivir) |  | References |
| --- | --- | --- | --- | --- | --- | --- |
|  |  |  |  | Vehicle | Remdesivir |  |
| attachment and entry | camostat mesilate | pancreatitis | Phase 2 | about 1 log reduction in an ex vivo model (PCLS) |  | (1, 2) |
|  | umifenovir | influenza | No clinical benefit | no in vivo data |  | (3) |
|  | baricitinib | rheumatoid arthritis | Phase 3 | no effect on virus replication |  | (4, 5) |
| initiation translation of polyproteins | Plitidepsin | multiple myeloma | Phase 2/3 | 1.5-2 log reduction, 4/8 achieved clearance in a mouse model | non-superior | (6) |
|  | Ternatin-4 | Preclinical compound |  | no in vivo data |  | (7, 8) |
|  | zotatifin | Phase 1 for cancer | Phase 1 | no in vivo data |  | (7, 8) |
|  | homoharringtonine | leukemia |  | 6/6 mice achieved clearance on 5 dpi |  | This study |
| proteolytic processing | MI-09 | SARS-CoV-2 | animal model | about 2 log reduction, clearance not achieved |  | (9) |
|  | MI-30 | SARS-CoV-2 | animal model | about 2 log reduction, clearance not achieved |  | (9) |
|  | lopinavir | AIDS | No clinical benefit | no effect on virus replication |  | (10) |
|  | ritonavir | AIDS | No clinical benefit | no effect on virus replication |  | (10) |
| transcription & RNA replication | remdesivir | coronavirus | FDA-approved | 10/36 rhesus macaques achieved clearance on 7 dpi |  | (11) |
|  | EIDD-2801/MK-4482 | coronavirus | Phase 2a | 5 log reduction assayed by PFU, 4/8 achieved clearance in the LoM model |  | (12) |
| multiple steps | clofazimine | leprosy | Phase 2 | 1-2 log reduction in hamster model, clearance not achieved | less-effective | (13) |
|  | ranitidine bismuth citrate | anti-ulcer |  | 1-1.5 log reduction, clearance not achieved | non-superior | (14) |

**Supplementary Table 2.** Homoharringtonine (HHT) exhibits broad-spectrum inhibition efficacy against coronaviruses.

| coronavirus | Abbreviation | Genus | IC50 (nM) | Reference |
| --- | --- | --- | --- | --- |
| Middle East respiratory syndrome coronavirus | MERS | Beta | 71.8 | Dyall et al. 2014( 15) |
| Mouse hepatitis coronavirus | MHV | Beta | 12 | Cao et al. 2015( 16) |
| Bovine coronavirus | BCoV | Beta | <<1uM | Cao et al. 2015( 16) |
| Human enteric coronavirus | HECoV | Beta | <<1uM | Cao et al. 2015( 16) |
| Porcine epidemic diarrhea virus | PEDV | Alpha | 112 | Dong et al. 2018( 17) |
| Severe acute respiratory syndrome coronavirus 2 | SARS-CoV-2 | Beta | ~100 | Wen et al. 2021( 18) |
| Severe acute respiratory syndrome coronavirus 2 | SARS-CoV-2 | Beta | 30 | lanevski et al. 2020( 19) |
| Severe acute respiratory syndrome coronavirus 2 | SARS-CoV-2 | Beta | 2.1uM** | Choy et al. 2020(20) |
| Porcine epidemic diarrhea virus | PEDV | Alpha | <<100 | This study |
| Porcine deltacoronavirus | PDCoV | Delta | <<100 | This study |
| Swine acute diarrhea syndrome coronavirus | SADS-CoV | Alpha | <100 | This study |

\*\* [Note - Choy et al.'s study reported a much larger IC50 value for multiple drugs, including HHT and Remdisivir (In Choy et al.'s study, IC50 of Remdisivir against SARS-CoV-2 was 26.9uM, while in many other studies, it was about 1uM(3, 21)).]

